# Epigenetic inheritance of telomere length in wild birds

**DOI:** 10.1101/284208

**Authors:** Christina Bauch, Jelle J. Boonekamp, Peter Korsten, Ellis Mulder, Simon Verhulst

## Abstract

Telomere length (TL) predicts health and lifespan in humans and other organisms, making the identification of the causes of TL variation of interest. At conception, zygotes inherit genes that regulate TL during early development, but at the same time already express a phenotype, which is the TL of the parental gametes that formed the zygote. Whether the effect of gamete TL is transient or affects TL for life depends on the extent to which regulatory genes compensate for gamete TL variation during early development. A carry-over effect of parental TL, resulting in epigenetic inheritance, has been suggested to explain the observed relationship between parental age and offspring TL in humans and other species. However, reports of parental age effects are based on cross-sectional data, and age at reproduction has numerous confounds. Furthermore, parental age may affect offspring telomere dynamics between conception and sampling, which could also explain the paternal age effect. Using longitudinal telomere data of jackdaw parents and their chicks, we show that chicks hatched with shorter telomeres as individual fathers aged, whereas mother age had no effect. By cross-fostering eggs, we confirmed the paternal age effect to be independent of paternal care after conception. The epigenetic effect accounted for 34% of the variance in offspring TL that was explained by paternal telomere length; the remaining 66% we ascribe to a combination of environmental and additive genetic effects. Thus, our results strongly indicate epigenetic inheritance of TL, with potential consequences for offspring fitness prospects.

**Significance statement:** Telomeres are DNA-protein structures at chromosome ends and their length predicts remaining lifespan in humans and other organisms. Variation in telomere length is thought to be largely of genetic origin, but telomere inheritance may be unusual because a fertilised cell already has a telomere length (most traits are first expressed later in life). Telomeres shorten with age, and, using long-term individual-based data of jackdaw families, we show that as fathers aged, they produced chicks with shorter telomeres. This shows that paternal telomere length directly affects offspring telomere length, i.e. is inherited genetically but without the involvement of genes. This is known as an epigenetic effect and explained a large part (≥34%) of the telomere resemblance between fathers and their offspring.

## INTRODUCTION

Telomeres are evolutionarily conserved DNA sequence repeats which form the ends of chromosomes together with associated proteins and contribute to genome stability (1). Telomeres shorten with age, at different rates in different tissues, depending on their rate of cell division (2) and on the activity of telomerase, a telomere elongating ribonucleoprotein (3). Age-adjusted telomere length (TL) relates to ageing-associated disorders and lifespan in humans (4, 5) and other organisms (6, 7). Given this relationship of telomeres with health and lifespan it is of importance to understand how variation in TL arises, which is already present early in life (8-10). TL heritability estimates are highly variable (11), but inheritance of TL is unusual compared with other traits, in that the phenotype is already expressed in the zygote without any effect of its own genome, because the zygote’s chromosomes have the telomeres of the two parental gametes. Subsequently, during development of the embryo, different telomere maintenance and restoration processes, under the control of different genes, regulate TL (12-16). In the course of development, these processes can potentially compensate fully for gamete derived differences in TL, in which case the effect of gamete TL is transient (Fig.1A). Alternatively, differences in gamete TL are carried over to later life (Fig. 1B). The latter case would imply an epigenetic component in the inheritance of TL, i.e. the inheritance of TL through changes in parental DNA that do not involve sequence changes (17, 18).

**Fig. 1.**
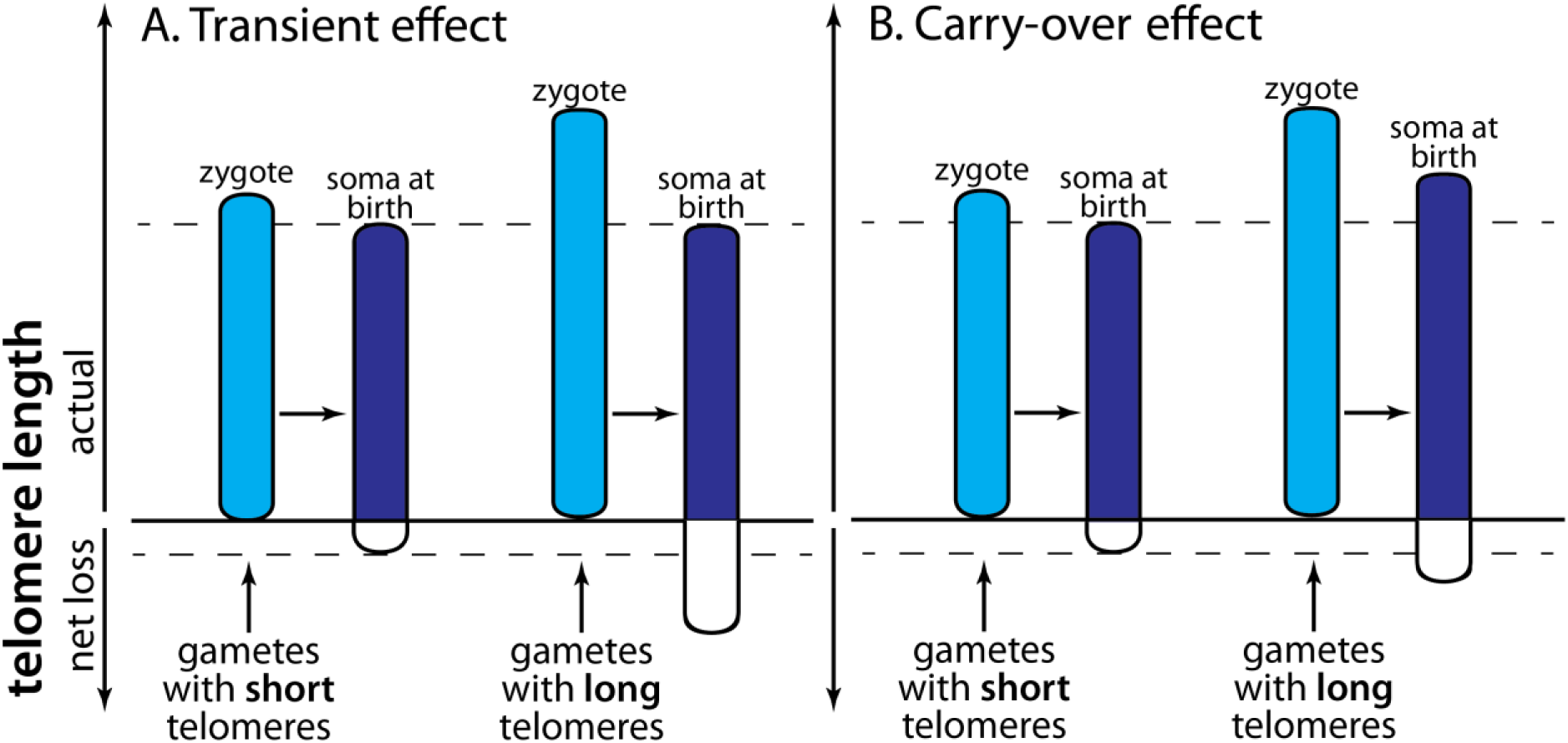
Schematic representation of inheritance of telomere length (TL). Shown is the TL of two individuals that differ in TL at conception due to a TL difference received via the gametes. Such a difference can arise through chance processes during gamete formation. TL shortens between conception and birth (2) and net TL loss is the outcome of cell division related shortening and telomerase-based restoration processes (3). Variation in genes regulating these processes cause the genetic inheritance of TL. We assume higher telomere attrition for the individual originating from gametes with long telomeres, because empirical evidence shows telomere attrition to be faster in individuals with longer telomeres (49), and because feedback mechanisms regulating TL are unlikely to achieve full compensation for TL differences at conception. If full compensation were achieved, this would prevent persistent epigenetic inheritance of TL and result in a transient effect (A) in the sense that TL at birth is independent of variation in gamete TL. If variation in gamete TL persist in later life, e.g. is still present at birth independent of genes that regulate TL during development, this implies an epigenetic carry-over effect (B).

Strongest evidence for epigenetic inheritance of TL comes from studies that show a relationship between parental (usually paternal) age and offspring TL (19). In humans, this trend parallels a qualitatively similar change in sperm TL with age, which is assumed to underlie the TL increase in offspring (20). However, studies of parental age effects in other species show mixed results and trends differ in direction between and within taxa (19, 21). More importantly, some critical uncertainties remain unresolved in any species. Firstly, studies to date are all cross-sectional, thus, comparing offspring of different parents that reproduced at different ages. Cross-sectional trends may differ from age related changes within individual parents when individuals with long TL are more likely to reproduce at older ages, which is not unlikely given the positive correlation between human TL and reproductive lifespan (22, 23). Secondly, parental age effects on offspring TL may arise from effects of parental age on developmental conditions prior to sampling. Because telomere attrition is highest early in life (2, 24), these effects can be substantial, as illustrated by parental age-effects on TL dynamics during development in European shags *Phalacrocorax aristotelis* (25) and Alpine swifts *Apus melba* (26). Lastly, due to their cross-sectional character, studies to date could not investigate whether changes in parental TL with age were predictive of changes in offspring TL with parental age at conception. These issues need to be resolved to establish whether the correlations between parental age and offspring TL can be attributed to epigenetic inheritance of TL, and before we can begin to understand why parental age effects on offspring TL appear to differ between and within taxa (19, 21).

To establish whether offspring TL changes with parental age at conception over the life of individual parents we used our long-term, individual-based dataset of free-living jackdaws *Corvus monedula*. Telomere data (measured in nucleated erythrocytes) from multiple chicks of the same parents that hatched up to 9 years apart allowed us to compare offspring TL within parents over time. As telomere attrition is highest early in life, we took blood samples for telomere analysis soon after hatching, when the oldest chick in a brood was 4 days old. To test if TL was influenced by age-dependent parental care prior to sampling, we cross-fostered clutches between nests, and tested whether foster parent age affected offspring TL. To investigate if the rate of telomere attrition within parents predicts the change in TL of the offspring they produce over consecutive years, we measured TL over the life of parents.

## RESULTS

### Parental age and offspring telomere length

As fathers aged, they produced offspring with 56±20 bp shorter TL for each additional year (father delta age effect in Table 1A, Fig. 2), thus showing that offspring TL declined with paternal age at conception within individual males. In contrast, there was no effect of maternal age on chicks’ TLs (Table 1B), neither when compared cross-sectionally, between offspring of different mothers over age (mean age mother in Table 1B), nor within mothers as they age (delta age mother). The negative, non-significant effect of maternal age on offspring telomere length is likely to be due to the age of their mates, because pair bonds in jackdaws are maintained over many years (pers. obs.) and hence maternal and paternal age are strongly correlated. This is confirmed by the finding that the observed delta age estimate in mothers is close to what would be expected based on the estimate found in fathers and the correlation of r=0.75 (n=298) between maternal and paternal age (0.75 * 56 bp = 42 bp, close to the estimate±SEM for delta mother age we observed, which was 38±23 bp; Table 1). The decline in offspring TL with father’s age was lower than the rate of TL attrition in the fathers themselves (−56±20 versus −87±15 bp/year). The variance in telomere attrition slopes between individual fathers was neither significant in the males themselves (p=0.09) nor in their offspring (p=0.14).

**Table 1.**
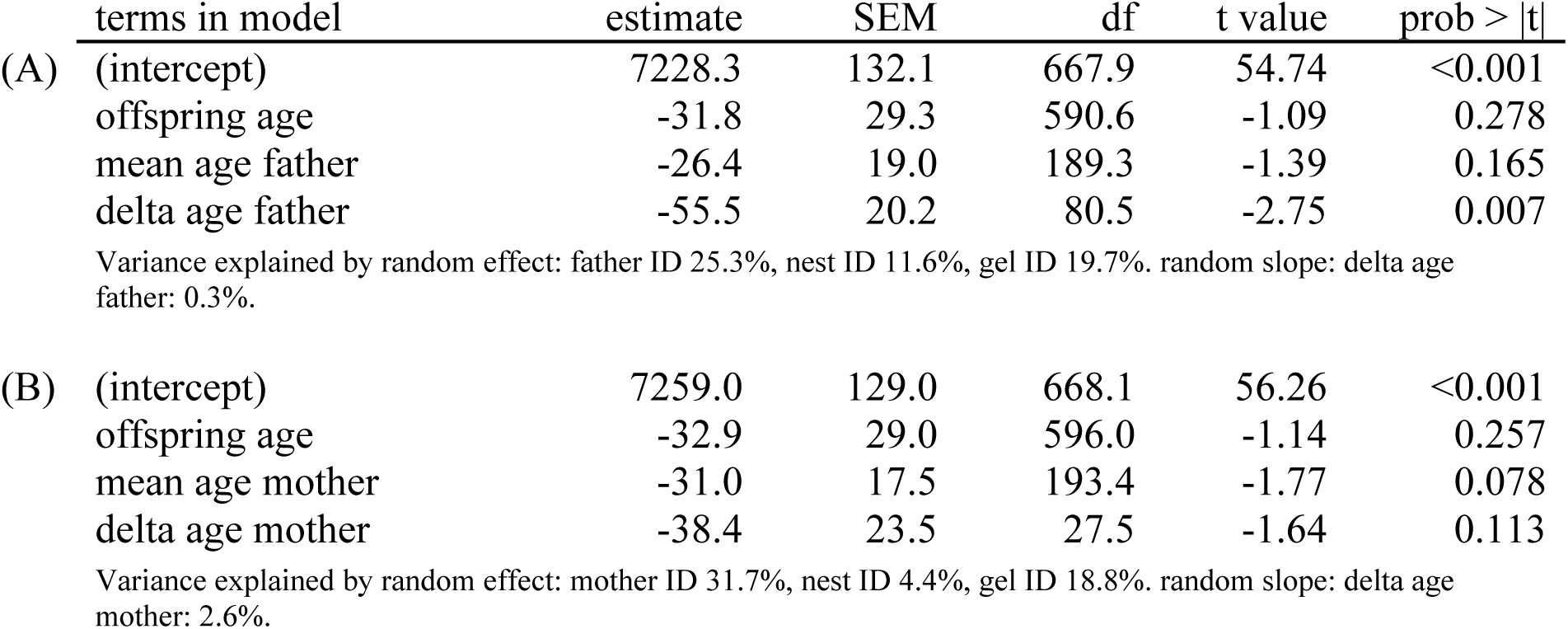
Linear mixed effects model to test the effect of parental age (years) on early-life telomere length of chicks in bp (n=715) – a comparison between (mean age parent) and within (delta age parent) fathers (A) and mothers (B). Model fit for (A) R^2^_GLMM(m)_=0.016, R^2^_GLMM(c)_=0.578 and (B) R^2^_GLMM(m)_=0.015, R^2^_GLMM(c)_=0.585. See also figure 2.

**Fig. 2.**
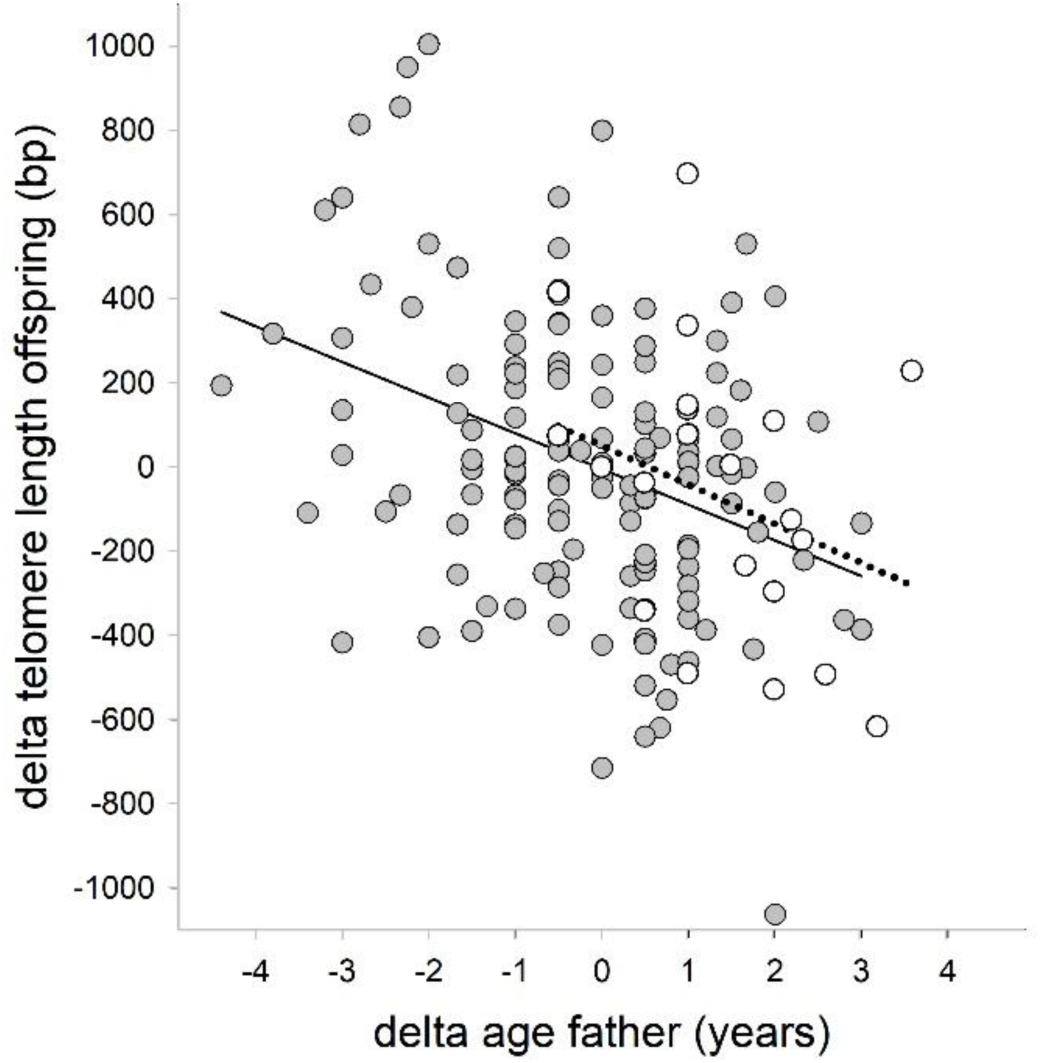
Offspring TL in relation to father age (delta offspring TL, i.e. TL per brood mean-centered per father; delta age, i.e. age mean-centered per father). Closed symbols and solid regression line for offspring from non-cross-fostered clutches, open symbols and dotted regression line for TL of offspring from cross-fostered clutches. As cross-fostering has been performed only in recent years, open symbols are not equally distributed over the x-axis.

### Test for early-life parental age effects on offspring TL

The paternal age effect on offspring TL could potentially be explained by age-dependent paternal care prior to sampling, when this affects telomere dynamics between conception and the age of 4 days. We tested this hypothesis by exchanging eggs between pairs shortly after clutch completion. Our analysis is based on telomere data of 61 chicks that hatched from 31 cross-fostered clutches. The coefficient of the age of the father caring for the eggs and offspring (i.e. the genetic father if not cross-fostered) and the coefficient of the age difference between genetic and foster father (which is 0 in case of no cross-fostering or age-match) were both negative and very similar (Table 2, Fig. 2). While the coefficient of the age difference did not quite reach statistical significance in a two-tailed test (p<0.09), we consider the similarity of the coefficients (10% difference) the more salient result. Thus, the older the father, the shorter the TL of his offspring, independent of the age of the male that cares for the eggs and offspring up to sampling. These results show that the paternal age effect on offspring TL is explained by the age of the genetic father and that the influence of the foster father age on offspring TL at age 4 days is negligible.

**Table 2.** Linear mixed effects model to test for early-life parental care effects associated with father age (years) on offspring telomere length (bp). Comparison of the effect of father age (for cross-fostered offspring foster father age) and the difference of genetic father age and

| terms in model | estimate | SEM | df | t value | prob > \|t\| |
| --- | --- | --- | --- | --- | --- |
| (intercept) | 7265.2 | 136.0 | 95.5 | 53.41 | <0.001 |
| offspring age | -34.9 | 29.4 | 581.1 | -1.19 | 0.236 |
| genetic or foster father age | -43.4 | 15.2 | 299.1 | -2.85 | 0.005 |
| delta father ages | -47.9 | 28.0 | 216.3 | -1.71 | 0.089 |

Variance explained by random effect: father ID 21.5.0%, nest ID 13.2%, gel ID 17.8%, analysis year 5.2%.
foster father age (delta father ages). $R^2_{\text{GLMM(m)}}=0.021$ , $R^2_{\text{GLMM(c)}}=0.578$ .

### Father telomere length and offspring telomere length

The paternal age effect on offspring TL raises the question whether the change in paternal TL with age predicts the change in offspring initial TL over years. We tested this by replacing mean age and delta age by mean and delta TL of the father (in the blood) in the year the offspring hatched. Fathers’ mean TL as well as delta TL were strongly and positively correlated with offspring TL (Table 3, Fig. 3). The effect of father’s mean TL on offspring TL can be attributed to additive genetic effects, possibly augmented by effects of a shared environment (11). The effect of fathers’ delta TL on offspring TL cannot be attributed to a genetic effect, because delta TL refers to variation within fathers over time. We therefore consider an epigenetic effect the most likely explanation of the effect of father’s delta TL on offspring TL. The variance explained by mean and delta paternal TL was 1.87 and 0.96 respectively, indicating that 34% (0.96 / 2.83) of the variance explained paternal TL can be attributed to the epigenetic effect. The slope of the epigenetic effect was 0.86±0.35, which is slightly higher than the 0.5 one would expect given that fathers contribute 50% of the zygote’s chromosomes, but not significantly so.

**Table 3.**
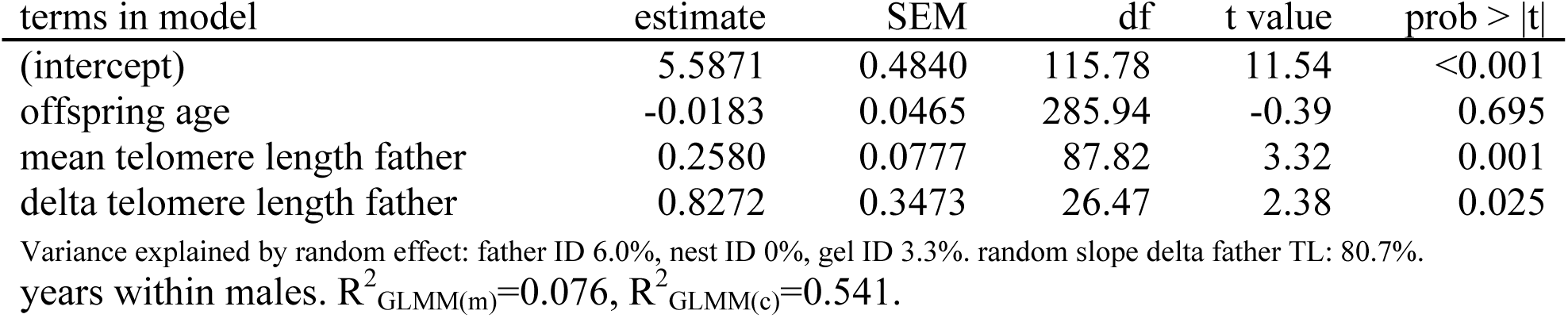
Linear mixed effects model to test the effects of father telomere length (kb) on their offspring telomere length (kb). Note that father TL variation was split in two components: the mean TL over all measurements per male (‘mean telomere length father’), and the deviation from that mean in each of the years he produced offspring sampled that year (‘delta telomere length father’. The coefficient of mean TL provides information on the effect of between make variation, while the coefficient of delta TL provides information on variation over the

| terms in model | estimate | SEM | df | t value | prob > \|t\| |
| --- | --- | --- | --- | --- | --- |
| (intercept) | 5.5871 | 0.4840 | 115.78 | 11.54 | <0.001 |
| offspring age | -0.0183 | 0.0465 | 285.94 | -0.39 | 0.695 |
| mean telomere length father | 0.2580 | 0.0777 | 87.82 | 3.32 | 0.001 |
| delta telomere length father | 0.8272 | 0.3473 | 26.47 | 2.38 | 0.025 |

**Fig. 3.**
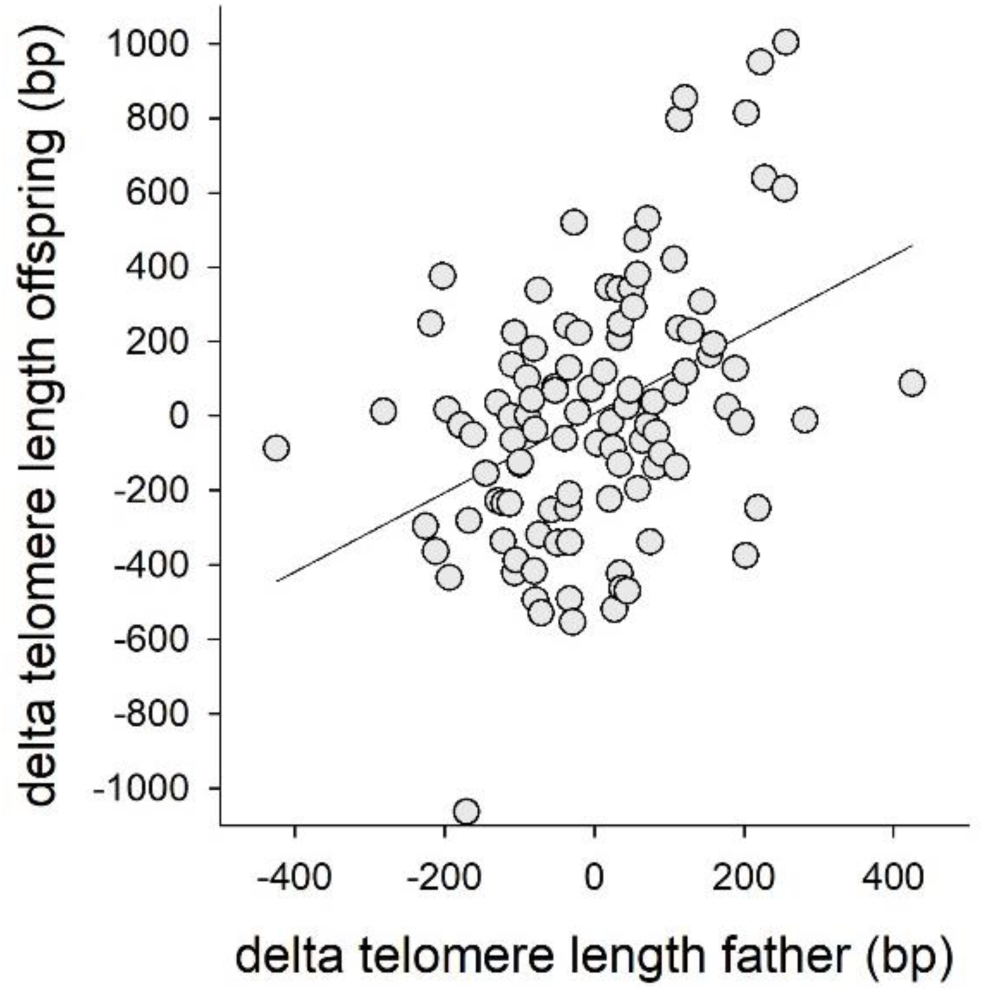
Offspring TL in relation to within individual variation in TL of their fathers at conception. Data on both axes are expressed relative to the mean TL of fathers (X-axis) or their offspring (Y-axis) over the different sampling years.

## DISCUSSION

Resemblance of TL between parents and offspring is potentially due to a dual inheritance mechanism, with on the one hand a ‘classic’ genetic effect through different alleles regulating TL during early development, and on the other hand an epigenetic effect of variation in TL in the gametes that at least in part carries through to later life (Fig.1). Suggestive evidence for an epigenetic contribution to the inheritance of TL comes from studies on humans showing a paternal age effect on offspring TL, but the available evidence is observational and cross-sectional (18, 19). We here show for the first time that offspring TL changed with age within individual fathers (i.e. longitudinally). Through a cross-foster experiment we confirmed that the paternal age effect cannot be attributed to age-dependent paternal care prior to sampling. From these findings, we infer that there is an epigenetic component in TL inheritance, which is further supported by the result that paternal, but not maternal age affected offspring TL. This is as expected, because sperm cells are formed throughout reproductive life, while the female’s complete stock of oocytes is formed before birth, and hence TL in female gametes is less prone to change (27-29). The negative relationship between paternal age and offspring TL is in accordance with results from a cross-sectional study in Alpine swifts (26), whereas cross-sectional studies on other bird species did not detect a paternal age effect (25, 30). However, the latter does not disagree with our findings, because also in our data set there was no significant cross-sectional association between paternal age and offspring TL (Table 1A).

While the longitudinal paternal age effect (represented by the variable ‘delta age father’ in Table 1A) in our study provides evidence for an epigenetic effect on offspring TL, we also found evidence suggesting an additive genetic effect when we examined the association between paternal and offspring TL directly. In this analysis, we separated between and within individual variation in paternal TL, and the between male component (represented by the variable ‘mean TL father’ in Table 3) shows the putative additive genetic effect on offspring TL, while the within male component (delta TL father) shows the epigenetic effect over the years within males. The coefficient of the between male effect was 0.26±0.08, suggesting the narrow sense heritability of jackdaw TL to be 0.52, which is well in the range observed in humans and other vertebrates (11). However, the variance in paternal TL between males is not entirely of genetic origin, because we show that in addition there is an epigenetic contribution to the variance. Thus, also the effect of ‘mean TL father’ in Table 3 can to an unknown extent be attributed to an epigenetic effect. Of the variance in offspring TL that was explained by paternal TL, we attributed approximately one third to epigenetic inheritance, but this ignores the epigenetic component in paternal TL, and hence the combined epigenetic effect will be more than one third of the resemblance between father and offspring TL.

Narrow sense heritability of human TL has been estimated using monozygotic and dizygotic twins (e.g. 31), assuming that a weaker resemblance between dizygotic twins compared to monozygotic twins can be attributed to the difference in relatedness. However, monozygotic twins develop from a single zygote, and hence from a single sperm cell and oocyte, and hence the difference in resemblance of a monozygotic versus a dizygotic twin pair may in part be due to an epigenetic effect of having developed from the same gametes (32). This process would lead to an overestimation of the narrow sense heritability compared to techniques that do not depend on twins.

The age effect on TL of the fathers themselves was almost twice as high as the paternal age effect on TL of their offspring. A two-fold difference suggests that blood and sperm TL shortened at the same rate, given that fathers contribute 50% of the zygote’s chromosomes. However, jackdaws reproduce for a few weeks in spring only, and presumably only produce sperm during this short period, because gonads in songbirds typically regress after the breeding season (33). Thus, the attrition rate of jackdaw sperm TL, which we assume to underlie the TL decline between offspring, is probably high during the short period of sperm production and low the rest of the year. That the rate of TL shortening in blood and sperm appears approximately equal may therefore well be a coincidence.

The direction of the paternal age effect in jackdaws (decreasing) is opposite to the direction of the paternal age effect in humans (increasing; 19). Assuming that paternal age effects in mammals (humans, chimpanzees; 19) and birds (jackdaws, alpine swifts; this study, 26) all reflect paternal age effects on sperm TL, this raises the question why these age effects on sperm TL are in the opposite direction. Seasonality of reproduction may well play a role, with species that produce sperm for a small part of the year having less need to maintain TL than species with year-round sperm production (19). The lengthening of TL in human sperm with age has been interpreted as the result of an overshoot in telomere maintenance (20) that can be interpreted as a safety margin in the maintenance process. Such a safety margin can be expected to be larger when the rate of sperm production and hence telomere attrition is higher. This may explain why chimpanzees, with a higher sperm production rate than humans (34), due to their promiscuous mating system, showed a steeper paternal age effect on offspring TL compared to humans (35). Information on the sign of the relation between paternal age and offspring TL in strongly seasonal mammal species and / or continuously reproducing bird species would allow a test of this hypothesis.

The epigenetic inheritance of TL potentially has more general implications. Parental age at conception has previously been shown to have negative effects on offspring fitness prospects in very diverse taxa, a phenomenon known as the *Lansing effect* (36-39). The underlying mechanisms are likely to be diverse but given that TL predicts survival in adult jackdaws (24), and TL early in life correlates strongly with TL in adulthood (8), offspring born to older fathers may have a shorter life expectancy due to their epigenetically induced shorter TL. A further aspect to explore is that the paternal age effect was previously found to lead to cumulative changes in TL over multiple generations (40). This could lead to population level changes in TL when the age structure of populations change, as has for example been observed in response to urbanisation (41). A population level change in TL may in itself have further demographic consequences (42), providing a positive or negative feedback, depending on whether increasing paternal age has a positive or negative effect on offspring TL.

## MATERIALS AND METHODS

### Data and blood sample collection

Life-history data and blood samples originate from an individual-based long-term project on free-living jackdaws *Corvus monedula* breeding in nest boxes south of Groningen, the Netherlands (53.14° N, 6.64° E). Jackdaws produce one brood per year with mostly 4 or 5 chicks. They are philopatric breeders and socially monogamous with close to zero extra-pair paternity (43, 44). During the breeding seasons around the hatching date nest boxes were checked daily for chicks. Freshly hatched chicks were marked by clipping the tips of the toenails in specific combinations and therefore the exact ages of offspring were known. Between 2005 and 2016, 715 jackdaw chicks were blood sampled at day 5 of the brood, i.e. as chicks hatch asynchronously, when the oldest chick(s) is (are) 4 days (mean age 3.6 days). These chicks originated from 298 nests, of 197 different fathers (age 1-13 years), whereof 66 were blood sampled repeatedly over years as well as their chicks (max. difference of age between chicks 8 years; 48% (32) of males had more than one partner over life) and 194 different mothers (age 1-11 years), whereof 62 were repeatedly blood sampled (max. difference of age between chicks 9 years; 52% (32) of females had more than one partner). 61 chicks (that contributed telomere data) hatched from 31 cross-fostered nests, i.e. eggs were exchanged between nest boxes (equal clutch sizes and laying dates (or up to 1 day difference)) soon after clutch completion. 54 (89%) of those chicks were fostered by a father of different age. Jackdaws in this project are marked with a unique colour ring combination and a metal ring. Parents were identified by (camera) observation during incubation and additionally later during chick rearing when caught for blood sampling (by puncturing the *vena brachialis*). Unringed adults were caught, ringed and assigned a minimum age of 2, as this is the modal recruitment age of colony-hatched breeders. All jackdaws were of known sex (molecular sexing (45)).

### Telomere analysis

Blood samples were first stored in 2% EDTA buffer at 4-7 °C and within 3 weeks snap frozen in a 40 % glycerol buffer for permanent storage at −80 °C. Terminally located TLs were measured in DNA from erythrocytes performing telomere restriction fragment analysis under non-denaturing conditions (24). In brief, we removed the glycerol buffer, washed the cells and isolated DNA from 5 µl of erythrocytes using CHEF Genomic DNA Plug kit (Bio-Rad, Hercules, CA, USA). Cells in the agarose plugs were digested overnight with Proteinase K at 50 °C. Half of a plug per sample was restricted simultaneously with *Hind*III (60 U), *Hinf*I (30 and *Msp*I (60 U) for ∼18 h in NEB2 buffer (New England Biolabs Inc., Beverly, MA, USA). The restricted DNA was then separated by pulsed-field gel electrophoresis in a 0.8 % agarose gel (Pulsed Field Certified Agarose, Bio-Rad) at 14 °C for 24h, 3V/cm, initial switch time 0.5 s, final switch time 7.0 s. For size calibration, we added ^32^P-labelled size ladders (1kb DNA ladder, New England Biolabs Inc., Ipswich, MA, USA; DNA Molecular Weight Marker XV, Roche Diagnostics, Basel, Switzerland). Gels were dried (gel dryer, Bio-Rad, model 538) at room temperature and hybridized overnight at 37 °C with a ^32^P-endlabelled oligonucleotide (5’-CCCTAA-3’)_4_ that binds to the single-strand overhang of telomeres of non-denatured DNA. Subsequently, unbound oligonucleotides were removed by washing the gel for 30 min at 37 °C with 0.25x saline-sodium citrate buffer. The radioactive signal of the sample specific TL distribution was detected by a phosphor screen (MS, Perkin-Elmer Inc., Waltham, MA, USA), exposed overnight, and visualized using a phosphor imager (Cyclone Storage Phosphor System, Perkin-Elmer Inc.). We calculated average TL using IMAGEJ (v. 1.38x) as described by Salomons *et al.* (24). In short, for each sample the limit at the side of the short telomeres of the distribution was lane-specifically set at the point of the lowest signal (i.e. background intensity). The limit on the side of the long telomeres of the distribution was set lane-specifically where the signal dropped below Y, where Y is the sum of the background intensity plus 10 % of the difference between peak intensity and background intensity. We used the individual mean of the TL distribution for further analyses. Samples were run on 92 gels. Repeated samples of adults were run on the same gel, chicks were spread over different gels. The coefficient of variation of one control sample from a 30-day old jackdaw chick run on 26 gels was 6% and of one control sample of a goose, with a TL distribution in a similar range, run on 31 other gels was 7%. Within-individual coefficient of variation for samples on the same gel was 3% (8, subset of the current data set).

### Statistical analyses

The relationships between parental age or father TL and early-life TL of offspring were investigated in a linear mixed effects model framework using a restricted maximum-likelihood method (testing specific predictions). To be able to separately evaluate between-and within-individual patterns of parental age or father TL, we used within-subject centering (46). Thus, instead of father age, mother age or father TL per se we introduced the mean value per individual over (if available) multiple years and delta age or delta TL, the deviation from that mean, respectively. To account for (genetic and potential other) similarities in TL between offspring of the same father or mother, we included father ID or mother ID as random effect in the model. As the dataset contains also siblings from the same year, we additionally added a random effect nest ID as a nested term in father ID or mother ID to the model investigating paternal or maternal age effects on TL, respectively. The age of chicks at sampling differed slightly (2-4 days) and as TL shortens with age (8), we included their age (in days) as covariate. We added gel ID as random effect. Analyses were performed separately for fathers and mothers. As indicators of model fit we provide marginal and conditional R^2^ values (47).

The cross-foster experiment was designed to test for potential effects of paternal age on early-life telomere attrition between egg laying and sampling (age 2-4 days). In a linear mixed model with offspring TL as dependent variable, we included both the age of the father caring for the clutch after cross-fostering and the age difference between the genetic father and foster father as covariates (age genetic father-age foster father). When the paternal age effect is independent of age-dependent effects between conception and sampling, we predict the coefficients of paternal age and the age difference between genetic father and foster father to be indistinguishable. In contrast, when the paternal age effect is entirely due to age-dependent paternal effects after laying, the coefficient will be the same, but opposite in sign. In case of a mixture of the two effects, the coefficient will be intermediate. In this analysis we used all offspring, i.e. also those that were not cross-fostered, and further included genetic father ID, nest ID, gel ID and year of telomere analysis as random effects, and chick age at sampling as covariate.

Statistics were performed in R (v. 3.3.3, R Development Core Team 2017, packages lme4, lmerTest, MuMIn).

## ACKNOWLEDGMENTS

We thank Martijn Salomons and the late Cor Dijkstra and all students involved in data and blood sample collection in the field, as well as Erica Zuidersma and Mike van het Land, who contributed to the analyses of TLs. Data were collected under license of the animal experimentation committee of the University of Groningen. CB was supported by a DFG research fellowship (BA 5422/1-1). JJB was funded by an NWO grant (823.01.009) awarded to SV.

